# Epidemiological dynamics of bacteriocin competition and antibiotic resistance

**DOI:** 10.1101/2020.10.23.351684

**Authors:** Sonja Lehtinen, Nicholas J. Croucher, François Blanquart, Christophe Fraser

## Abstract

Bacteriocins, toxic peptides involved in the competition between bacterial strains, are extremely diverse. Previous work on bacteriocin dynamics has highlighted the role of non-transitive ‘rock-paper-scissors’ competition in maintaining the coexistence of different bacteriocin profiles. The focus to date has primarily been on bacteriocin interactions at the within-host scale. Yet, in species such as *Streptococcus pneumoniae* with relatively short periods of colonisation and limited within-host diversity, ecological outcomes are also shaped by processes at the epidemiological (between-host) scale. Here, we first investigate bacteriocin dynamics and diversity in epidemiological models. We find that in these models, bacteriocin diversity is more readily maintained than in within-host models, and with more possible combinations of coexisting bacteriocin profiles. Indeed, maintenance of diversity in epidemiological models does not require rock-paper-scissors dynamics; it can also occur through a competition-colonisation trade-off. Second, we investigate the link between bacteriocin diversity and diversity at antibiotic resistance loci. Previous work has proposed that bacterial duration of colonisation modulates the fitness of antibiotic resistance. Due to their inhibitory effects, bacteriocins are a plausible candidate for playing a role in the duration of colonisation episodes. We extend the epidemiological model of bacteriocin dynamics to incorporate an antibiotic resistance locus and demonstrate that bacteriocin diversity can indeed maintain the coexistence of antibiotic sensitive and resistant strains.

## Introduction

Bacteriocins are toxic peptides that allow bacteria to eliminate competitors. Bacteriocins systems are pervasive in bacterial species and are thought to play a significant role in competition within (and possibly between [1]) species [2]. A mechanistic understanding of the role of bacteriocins in competition is therefore important for characterising the ecological dynamics of bacteria.

At a high level of abstraction, bacteriocin systems can be thought of as consisting of three components: a system to produce and secrete toxins, a system to achieve immunity against these toxins, and a regulatory system to control toxin and immunity production—which may involve release of signalling molecules (‘pheromones’) allowing quorum-sensing and kin recognition. Even at this level of abstraction, bacteriocin systems are highly diverse. A single species may have a number of distinct systems, with considerable diversity within each of these: coexistence of strains with different combinations of toxin, immunity and regulatory genes is pervasive [3].

For example, the bacterial species *Streptococcus pneumoniae* has multiple different bacteriocins systems. The two best described systems—the *blp* (bacteriocin-like peptide) locus [4, 5, 6, 7] and the *cib* (competence-induced bacteriocin) locus [8]—are both ubiquitous. The specific combination of genes that make up the *blp* locus is highly variable between strains, while the *cib* locus is more conserved. Both loci are associated with a pheromone signalling system, which regulates the expression of toxins and immunity. These signalling systems are also diverse: there are multiple distinct ‘pherotypes’, i.e. specific pheromone-receptor pairs allowing targeted signalling to cells of the same pherotype [9, 10]. In contrast, other pneumococcal bacteriocin systems, such as the *pld* locus [11] and the circular toxin pneumocyclicin [12] are found on only a subset of strains. Thus, to fully capture bacteriocin competition, models should be able to explain both the diversity in bacteriocin profiles and the variation in the specifics of this diversity between different bacteriocin systems.

Previous theoretical and experimental work has highlighted the role of non-transitive competition in maintaining bacteriocin diversity. Indeed, two-strain models of toxin-producing (‘producer’) and toxin-susceptible (‘non-producer’) strains do not predict coexistence: depending on the effectiveness of the toxins and the cost associated with their production, either the producer or the non-producer strategy out-competes the other [13, 14]. Coexistence can be achieved by inclusion of a third (‘immune’) strain which does not produce toxins but does have immunity against them (or, alternatively, produces less effective and less costly toxins [14]). If both toxin and immunity incur a fitness cost, the total cost to the producer strain (which has both toxins and immunity), is greater than the total cost to the immune strain (which only incurs the cost of immunity, but not toxin production). As a consequence, in head-to-head competition, the non-producer out-competes the immune strain; the immune strain out-competes the producer; and the producer out-competes the non-producer. This type of non-transitive competitive structure is referred to as ‘rock-paper-scissors’ dynamics [15].

To provide additional clarity in the context of bacteriocin dynamics, we make a distinction between two types of rock-paper-scissors model. In the most general formulation of rock-paper-scissors dynamics (Figure 1 A) [15, 16], strain interactions are considered in terms of head-to-head competition for the occupation and invasion of patches or sites. Strains are differentiated through their ability to invade occupied patches: each strain wins against one of the other strains and loses against the other. Apart from the outcomes of this head-to-head competition, there are no further ecological differences between the strains. Models building on this structure have been studied in a number of ecological contexts [15], including bacteriocin dynamics [16]—where, at the epidemiological scale, patches would represent hosts to colonise, or, at the within-host scale, space to occupy in the colonised niche. Such models predict oscillatory dynamics of the three strains in well-mixed environments and stable coexistence in spatially structured environments [15, 16].

**Figure 1:**
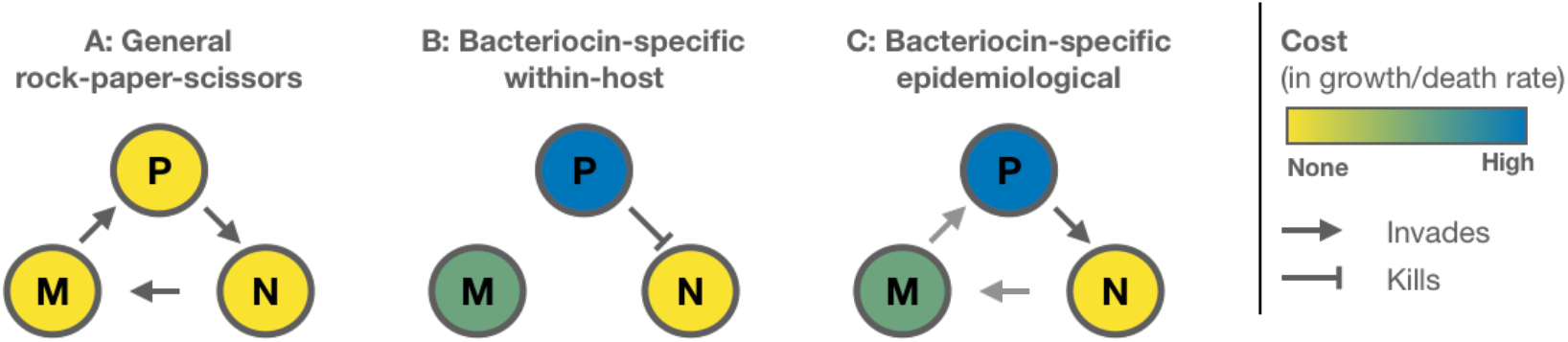
Strain interactions arising in different models of bacteriocin dynamics. *P* represents the producer strain (both toxins and immunity); *M* the immune strain (immunity only) and *N* the non-producer (no toxins, no immunity). A: A general formulation of rock-paper-scissors competition. Strain interactions are modelled in terms of the outcome of head-to-head competition between two strains. The costs of toxins and immunity, as well as the killing action of the toxins, are modelled implicitly in the outcome of this head-to-head competition. At the epidemiological scale, assuming fitness differences between strains are only apparent when the strains are competing for the same host gives rise to this model structure. B: Bacteriocin-specific within-host model [14, 17]. The producer strains kill non-producer strains, leading to resources being freed up. The costs of immunity and bacteriocin production are modelled as decreased growth rate or increased death rate. C: Bacteriocin-specific epidemiological model. The killing of the non-producer strain by the producer strain leads to the host becoming colonised with the producers strain. In this epidemiological model, the costs associated with bacteriocin production and immunity can be represented in terms invasion probabilities and/or epidemiological parameters (i.e. transmission and/or clearance rates).

This general formulation of rock-paper-scissors competition contrasts with bacteriocin-specific rock-paper-scissors models (Figure 1 B) [14, 17] that explicitly represent bacteriocin-related processes within a host (or within a liquid culture or Petri dish). There are two key differences between these models. The first is in the interaction between producer and non-producer strains: in the bacteriocin-specific formulation, toxins lead to the death of non-producer cells, rather than their direct replacement by the producer cells. The death of non-producer cells frees up resources (e.g. nutrients or space), which can then be used by any of the strains. As a result, the immune strain gains the same benefit from the production of toxins as the producer strain, without paying the cost (a ‘cheater’ strategy). The second difference is how the cost of immunity and bacteriocin production are represented: these are not modelled in terms of strain interactions, but rather as a reduced growth rate or increased death rate for the immune and producer strains. These bacteriocin-specific models predict stable coexistence of the three strains in spatially structured environments, but dominance of a single strain in well-mixed systems [14]. This prediction has been verified in an experimental model of producer, non-producer and immune strain interactions in *Escherichia coli* [17]. More recent modelling suggests that when the benefits of immunity are shared, rather than specific to the immune strain (e.g. immunity involves secretion of toxin-degrading compounds), coexistence of strains can also arise in well-mixed environments [18].

These bacteriocin-specific models capture within-host interactions between different bacteriocin profiles and thus provide insights into dynamics at this scale. Yet, within-host interactions will also impact dynamics at the epidemiological scale by allowing strains to invade already colonised hosts and/or by preventing such invasion. In addition, any fitness costs associated with bacteriocin production and immunity may also impact epidemiological parameters—i.e. transmission and clearance rates. In species such as *Streptococcus pneumoniae*, these effects at the epidemiological scale are likely to play an important role in shaping the ecology and evolution of bacteriocin systems, for two reasons. Firstly, within-host models assume all three species are co-colonising the same host. However, multiple colonisation is not necessarily the norm: for example, reported rates of co-colonisation with multiple pneumococcal strains are around 25% to 50% depending on setting [19, 20, 21]. Secondly, when the duration of colonisation is relatively short (e.g. of the order of months in *S. pneumoniae*), factors affecting transmission will play a large role in shaping allele frequencies in the overall bacterial population.

The impact of bacteriocins at the epidemiological scale is not well established and may depend on the specific bacteriocin system. For example, the *cib* locus provides established colonisers an advantage over invaders in a murine model of pneumococcal colonisation [22], suggesting these bacteriocins play a defensive role. On the other hand, an observational study of pneumococcal strains co-colonisation in humans suggest that the *blp* locus has a limited impact on which strains can co-colonise hosts [23]. Furthermore, the impact of the cost of bacteriocin production and immunity on epidemiological parameters and the effects of kin recognition are unclear. Exploring bacteriocin dynamics at the epidemiological scale therefore requires a flexible modelling approach that can capture a range of effects.

While the general rock-paper-scissor model (Figure 1A) can be interpreted as an epidemiological model (with patches corresponding to hosts), previous work has focused on patch invasion and not explored the possibility that toxin-production and immunity associated costs might impact epidemiological parameters. Bacteriocin-specific within-host models (Figure 1B), even with the inclusion of spatial structure, do not accurately capture interactions at the epidemiological scale. Although the presence of hosts structures the bacterial population, the interaction between the producer and non-producer is not equivalent at the two scales: at the within-host scale, toxins lead to the death of the non-producer and freeing up of resources, while at the epidemiological scale, toxins lead to invasion of the host occupied by the non-producer and no freeing up of hosts. Thus, bacteriocin-specific models at the epidemiological scale (Figure 1C) are necessary for understanding bacteriocin dynamics and diversity in *S. pneumoniae* and species with similar ecology.

A further motivation for developing epidemiological models of bacteriocin dynamics is their potential role in determining the duration of colonisation episodes. We have previously suggested that duration of colonisation affects selection pressure for antibiotic resistance in species which are carried asymptomatically most of the time, such as *S. pneumococcus* [24]. Strains that colonise hosts for longer gain a greater fitness benefit from resistance, leading to an association between long duration of colonisation and antibiotic resistance (including multi-drug resistance [25]). As a result, balancing selection maintaining diversity at a locus that affects duration of colonisation could also maintain diversity at a resistance locus [24], providing a potential explanation to the long-standing puzzle of why antibiotic resistance has not yet reached fixation [26]. However, this explanation is not complete, because the genetic determinants of duration of colonisation have not been fully identified [27]. A more complete understanding of resistance dynamics therefore requires identifying loci that contribute to variation in duration of colonisation. The role of toxin production in killing competing bacteria and the role of immunity in preventing this killing makes bacteriocin loci a promising candidate.

This paper is organised in two parts: the first part develops an epidemiological model of bacteriocin dynamics in *S. pneumoniae* (or a species with similar ecology). We explore the circumstances in which this model allows coexistence of strains with different bacteriocin profiles. In the second part, we investigate differences in duration of colonisation between bacteriocin profiles and their impact on resistance dynamics.

### Part I: Bacteriocin dynamics

#### Epidemiological modelling of bacteriocin dynamics

We begin by considering a bacteriocin system with two components: a toxin gene and an immunity gene. There are therefore three possible strains: a producer strain (*P*), with both toxin and immunity; an immune strain (*M*), with immunity but no toxin; and a non-producer strain (*N*), with neither toxin nor immunity (i.e. the entire bacteriocin locus is absent). Strains with the toxin but no immunity are assumed to be lethal to themselves and therefore not included. We formulate a generic epidemiological model of competition between three strains rather than focusing on what is known about a specific bacteriocin system, and consider how ecological differences between bacteriocin profiles can be reflected in this generic model.

In this model formulation, strain *i* colonises uncolonised individuals *X* at rate *β*_*i*_ and is cleared at rate *µ*_*i*_. In addition, strains can invade already colonised individuals and displace the resident strain at rate *β*_*i*_*k*_*ij*_ for strain *i* replacing strain *j*, where *k* represents the probability of successful invasion relative to colonisation of an uncolonised host. We assume the dynamics of this replacement are fast; the invading strain is therefore modelled as replacing the resident strain instantaneously. Hosts are therefore only ever colonised with a single strain at a time and there is no co-infection. Note that our qualitative results are robust to relaxing this assumption (SI Section 2.1). The dynamics of this general model are described by the following equations:

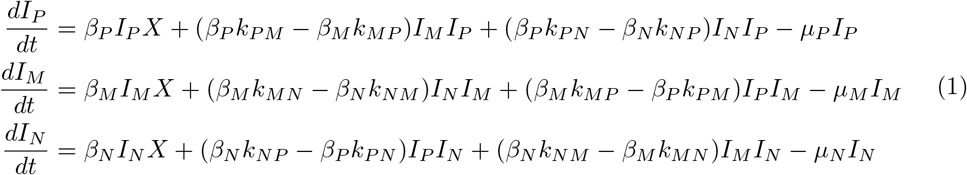

Here, the *X* and *I* variables represent proportions of the total host populations: *X* + *I*_*P*_ + *I*_*M*_ + *I*_*N*_ = 1.

Differences between bacteriocin profiles (i.e. producers *P*, non-producers *N* and immune strains *M*) can be reflected in two ways in this model (Figure 1 C). Firstly, through asymmetric invasion probabilities *k* (0 *≤ k ≤* 1) arising from differences in within-host fitness: toxins allow *P* to invade *N* more often than the other way around (*k*_*P N*_ *≥ k*_*NP*_); similarly, the cost of toxin production means *M* out-invades *P* (*k*_*MP*_ *≥ k*_*P M*_); and the cost of immunity means *N* out-invades *M* (*k*_*NM*_ *≥ k*_*MN*_). Secondly, the costs of toxins and immunity could also affect epidemiological parameters—i.e. transmission and clearance rates: *µ*_*P*_ *≥ µ*_*M*_ *≥ µ*_*N*_ and *β*_*N*_ *≥ β*_*M*_ *≥ β*_*P*_. These inequalities reflect typical assumptions about bacteriocin dynamics and, unless otherwise indicated, we have assumed they hold in our analysis.

In this structure, the impact of toxins (and other within-host fitness differences) can be modelled as either offensive (i.e. enabling invasion of already colonised hosts) or defensive (i.e. preventing invasion by another strain) or a combination of both. Figure 1 depicts offensive interactions: invasion is only possible when the invading strain has a fitness advantage. This is also how we parametrise the model in main text results (i.e. *k*_*NP*_ = *k*_*P M*_ = *k*_*NM*_ = 0). It is worth noting that the distinction between offensive and defensive only impacts epidemiological dynamics if the transmission rate differs between strains: when transmission rates are equal, the interaction between two strains—say strain *A* and strain *B*—simplifies to [*k*_*AB*_ *− k*_*BA*_]*βI*_*A*_*I*_*B*_ in the dynamics of strain *A* (and similarly [*k*_*BA*_ *− k*_*AB*_]*βI*_*A*_*I*_*B*_ in the dynamics of strain *B*). The dynamics therefore depend only on the *relative* rates of invasion. Under these circumstances, modelling bacteriocins as offensive is mathematically equivalent to modelling them as defensive. This equivalence does not hold when transmission rates are not equal (i.e. when epidemiological costs affect transmission rate). However, we find that in practice, the distinction between offensive and defensive bacteriocins does not have an impact on which strains are observed to coexist (SI Section 2.2).

#### Epidemiological differences between strains allow a wider range of outcomes

Our aim is to understand the circumstances under which multiple strains coexist at equilibrium in this system. We address this through linear stability analysis (unless otherwise indicated). In general, we parametrise clearance rate as *µ* = 1 and transmission rate as *β* = 3. In time units of month^-1^, these are plausible values for *S. pneumoniae* [26]. All analyses and simulations were performed using Wolfram Mathematica [28]; the code is available as a Supporting File.

First, it is useful to note that if strains differ only in invasion probabilities (the *k* parameters) but not epidemiological parameters (i.e. *µ*_*P*_ = *µ*_*M*_ = *µ*_*N*_ and *β*_*N*_ = *β*_*M*_ = *β*_*P*_), the model is structurally identical to the classic rock-paper-scissors model (e.g. [15], see Figure 1). The behaviour of this system is well-characterised: as long as the non-transitive competitive structure is maintained, all three strains coexist, with oscillatory dynamics around a stable equilibrium point (see also Figure 2). When the non-transitive competition structure is not present, a single strain dominates. Equilibria involving just two of the strains are never stable.

**Figure 2:**
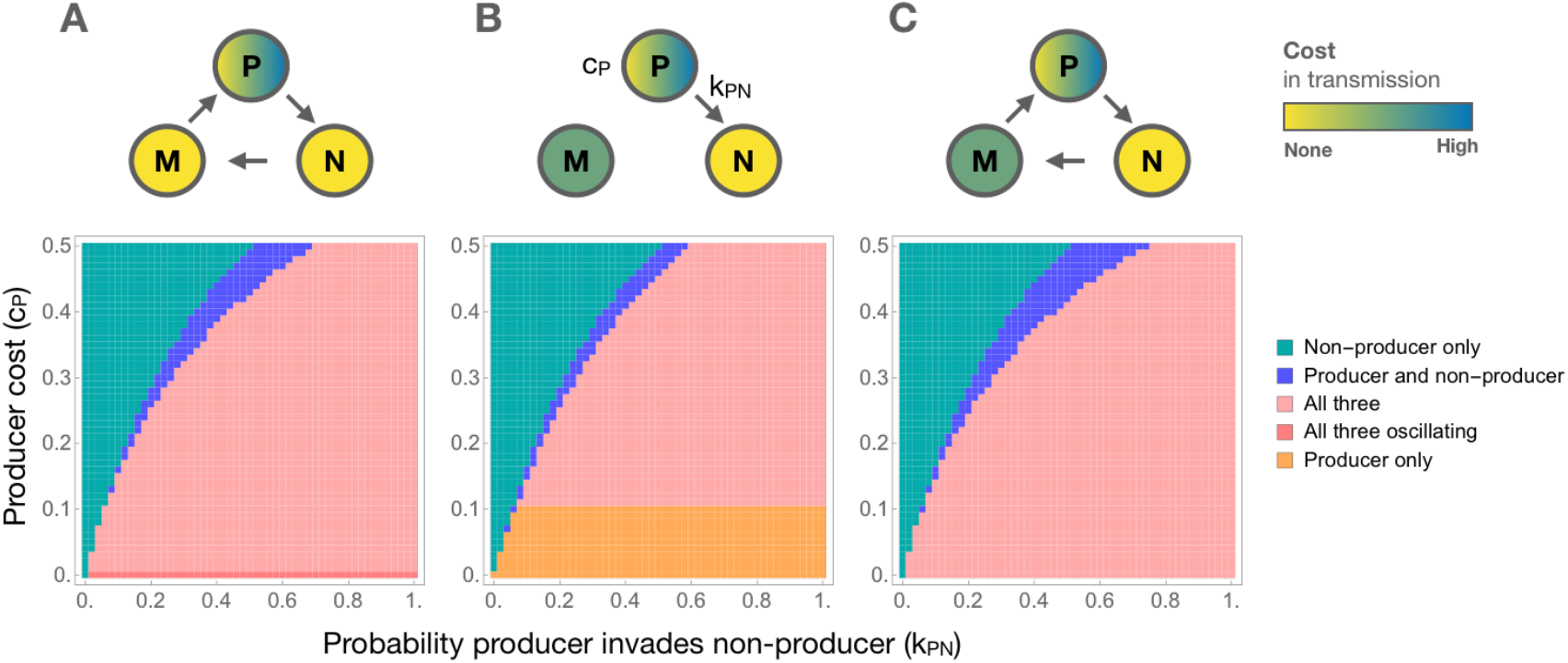
Outcomes of strain competition in an epidemiological model of bacteriocin competition for different parameter values. *P* represents the producer strain, *M* the immunse strain, and *N* the non-producer strain. The top panels represent the interactions between different bacteriocin profiles. For each of the three bottom panels, the x-axis represents the probability that the producer invades the non-producer (*k*_*P N*_); the y-axis represents the reduction in transmission rate resulting from the fitness cost incurred by the producer strain (*β*_*P*_ = *c*_*P*_ *β*); and the color indicates which strains are present at equilibrium. **A** Fitness differences between bacteriocin-profiles are modelled as a within-host competitive advantage, allowing strains to displace each other in colonisation (*k*_*MP*_ = *k*_*NM*_ = 0.5, *k*_*P N*_ *≥* 0 and all other *k* parameters are 0). This results in a non-transitive competitive structure. When the producer strain is associated with no additional cost in transmission, this model identical to the classic rock-paper-scissors model and predicts oscillatory dynamics. Introducing a cost in transmission removes the oscillatory dynamics and widens the possible range of outcomes. **B** Strain displacement is only possible through bacteriocin production (i.e. *k*_*P N*_ *≥* 0, all other *k* parameters are 0). The fitness costs incurred by the producer and the immune strain are modelled as a decreased transmission rate (the cost to the immune strain *c*_*M*_ is 0.1). When the producer strain has a smaller cost than the immune strain (implying an additional benefit to bacteriocin production - e.g. inhibitory action against other species), the producer strain excludes the other two strains. **C** A combination of the two previous panels: fitness differences between bacteriocin profiles affect both invasion probabilities and transmission (*c*_*M*_ = 0.1, *k*_*MP*_ = *k*_*NM*_ = 0.5, *k*_*P N*_ *≥* 0 and all other *k* parameters are 0). For all panels, *β* = 3 and *µ* = 1.

Introducing differences in epidemiological parameters—i.e. transmission rate *β* or clearance rate *µ*—between bacteriocin profiles has two effects. Firstly, oscillation is not observed when the strains differ in epidemiological parameters (Figure 2 and SI Figure S4). Thus, like spatial structure in previous models [14, 15, 17], differences in the epidemiological parameters of strains act to stabilise coexistence. Secondly, introducing differences in the range of possible outcomes increases. Stable coexistence no longer requires all three strains; for some parameter ranges, the producer and non-producer strain can coexist without the immune strain (Figure 2 B). Such two-strain coexistence is not observed in previous models of bacteriocin dynamics.

It is worth noting that the properties of the two and three strain equilibria are different. The general rock-paper-scissors model (Figure 1 A) is known to give rise to a ‘survival of the weakest’ effect: increasing the competitive advantage of a strain (i.e. ability to invade) *decreases* its equilibrium frequency [15]. This occurs because of the non-transitive competitive structure. For example, a smaller *k*_*P N*_ and thus less frequent displacement of *N* by *P* increases the frequency of *N* and therefore the rate at which *M* is displaced. This, in turn, decreases the frequency of *M* and displacement of *P*, leading to a higher frequency of *P* despite its lower competitive advantage. We find that this effect also applies to the relationship between transmission rate and equilibrium frequency (SI Figure S3 and SI Section 2.3): increasing the transmission rate of a strain decreases its equilibrium frequency when all three strains are present at equilibrium. This survival of the weakest effect is not observed for two-strain equilibria.

Finally, the number of stable strain combination is even greater if bacteriocin-production has other benefits beyond inter-strain competition—such as bacteriocidal action against other species leading to the producer strain also having increased success in colonising hosts which are not carrying the focal species. This would result in an increased transmission rate for the producer strain, and thus offset some of the cost of bacteriocin production. Under these circumstances, the producer strain can exclude the other two strains (Figure 2 B).

### The effects of kin recognition (pherotype)

#### Extension of models to include pherotype

To explore the effect of kin recognition on bacteriocin diversity, we expand the original model to include two different signalling molecules and corresponding receptors (i.e. two distinct pherotypes), yielding a six strain model. The dynamics between strains of the same pherotype remain as described in Equations 1. Across pherotypes, immunity is not effective: the signal to turn on expression of immunity proteins is not recognised. Producer strains are therefore able to out-compete immune strains with a different pherotype; in other words, the interaction between producer and immune strain is the same as the interaction between producer and non-producer strain. Under the assumption that bacteriocins are required for invasion, this results in the model depicted in Figure 3A (see SI Section 1.1 for equations and additional discussion).

**Figure 3:**
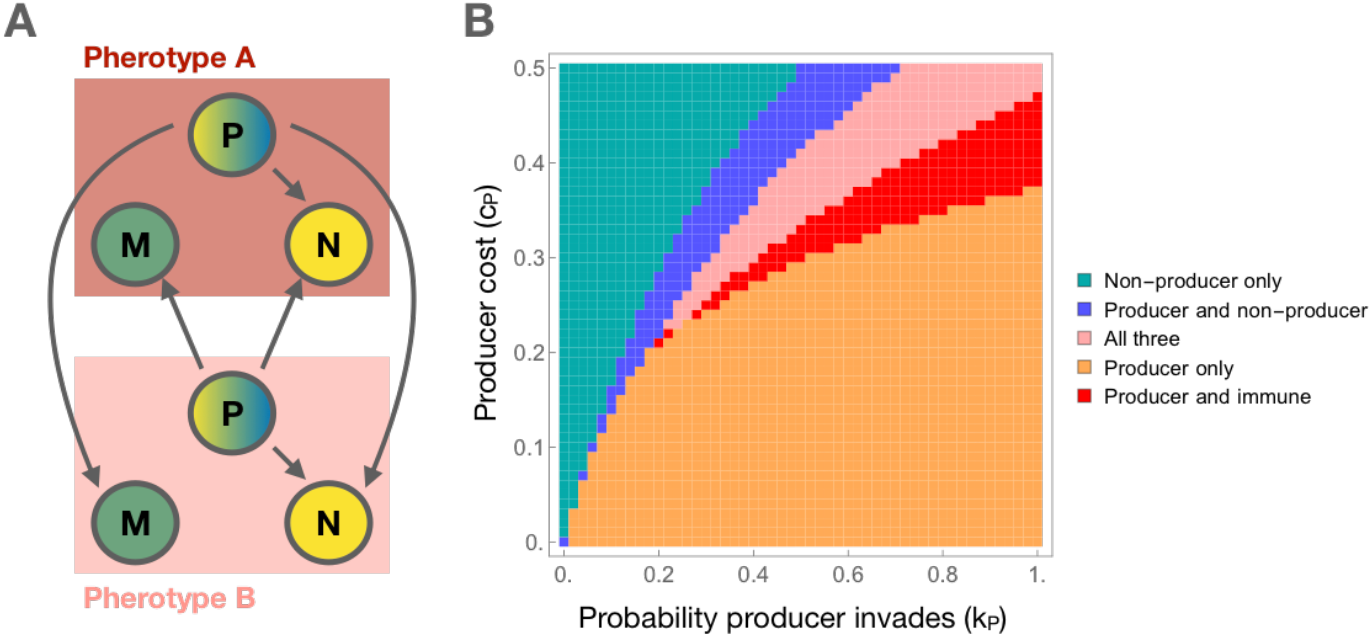
The effect of kin recognition on the coexistence of different bacteriocin profiles in an epidemiological model. **A** Strain interactions in the pherotype model. We assume bacteriocins are required for invasion (corresponding to Figure 2 B). Immunity is not effective across pherotypes, allowing the producer strain of pherotype A to displace the immune strain of pherotype B (and vice-versa). (See SI Section 1.1 for further discussion of possible strain interaction). The properties of pherotypes A and B are otherwise identical. **B** The inclusion of pherotype further increases possible outcomes but decreases the parameter space in which multiple strains coexist. As in Figure 2, the x-axis represents how well the producer strain invades (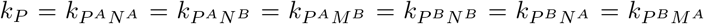, where the *A* and *B* superscripts indicate pherotype); the y-axis represents the transmission cost for the producer strain; and the colours indicate which strains are present at equilibrium. Other parameters are *β* = 3, *µ* = 1, *c*_*M*_ = 0.1 and all other *k* parameters 0, similar to Figure 2 B. Due to the difficulty of computing equilibrium solutions and eigenvalues for this model, these results are derived through simulating rather than stability analysis (simulation until *t* = 10^5^ months).

The inclusion of pherotype in the model decreases the parameter space in which coexistence is observed: the producer strain excludes the other strains in a large part of the explored parameter space (Figure 3B). The inclusion of pherotype also increases the number of combinations in which strains can coexist, allowing stable equilibria with the producer and immune strains without the non-producer strain. These effects arise because the addition of pherotype expands the range of strains that are susceptible to the toxins, thus allowing the producer strain to exist without the non-producer strain.

### Part II: Potential role for bacteriocins in resistance dynamics

#### Bacteriocins profiles differ in duration of colonisation

We now turn to the potential role of bacteriocins in the dynamics of antibiotic resistance. We have previously suggested that the duration of colonisation affects selection pressure for antibiotic resistance: modelling predicts that fitness effect of resistance depends on a strain’s duration of colonisation, with longer colonisation being associated with a greater benefit from resistance [24]. Indeed, a strain’s duration of colonisation correlates with the frequency of resistance within the strain in multiple pneumococcal datasets [24, 29]. Variation in the duration of colonisation among coexisting strains could therefore maintain the coexistence of antibiotic sensitive and resistant strains, with long-colonisers being antibiotic resistant and short-colonisers antibiotic sensitive. Bacteriocins, due to their inhibitory effects on others strains, are a plausible source of such stable variation in duration of colonisation.

To assess the potential role of bacteriocins in resistance dynamics, we examine the extent of variation in duration of colonisation between different bacteriocin profiles. In the model represented by Equations 1, the mean duration of colonisation of strain *i* (*D*_*i*_)—i.e. the inverse of its overall clearance rate—is given by:

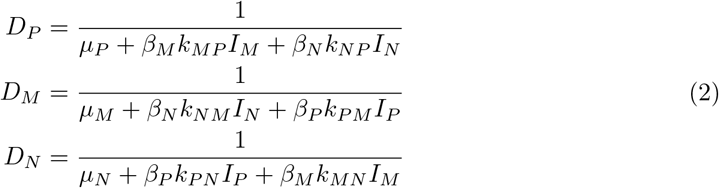

Bacteriocin profiles can thus affect duration of colonisation through two mechanisms: i) directly through any effects of the cost of toxin production and immunity on clearance rate and ii) through strain displacement, the impact of which depends on invasion rates (*k*), the effect of the cost of immunity and toxin production on transmission rate, and the prevalence of the invading strain. As a result, the extent of variation in duration of colonisation and which strain is associated with the longest colonisation depends on parameter values (Figure 4, and SI Section 2.5). When costs affect transmission, the longest duration of colonisation is associated with either the producer or immune strain, and the shortest with either the non-producer or immune strain. This is reversed when costs affect clearance, with the longest duration of colonisation associated with the non-producer or immune strain, and the shortest with the producer or immune strain. In broad terms, the greatest variation in duration of colonisation is observed when the cost associated with bacteriocin production is high. A more detailed discussion of the relationship between parameters and duration of colonisation can be found in SI Section 2.5.

**Figure 4:**
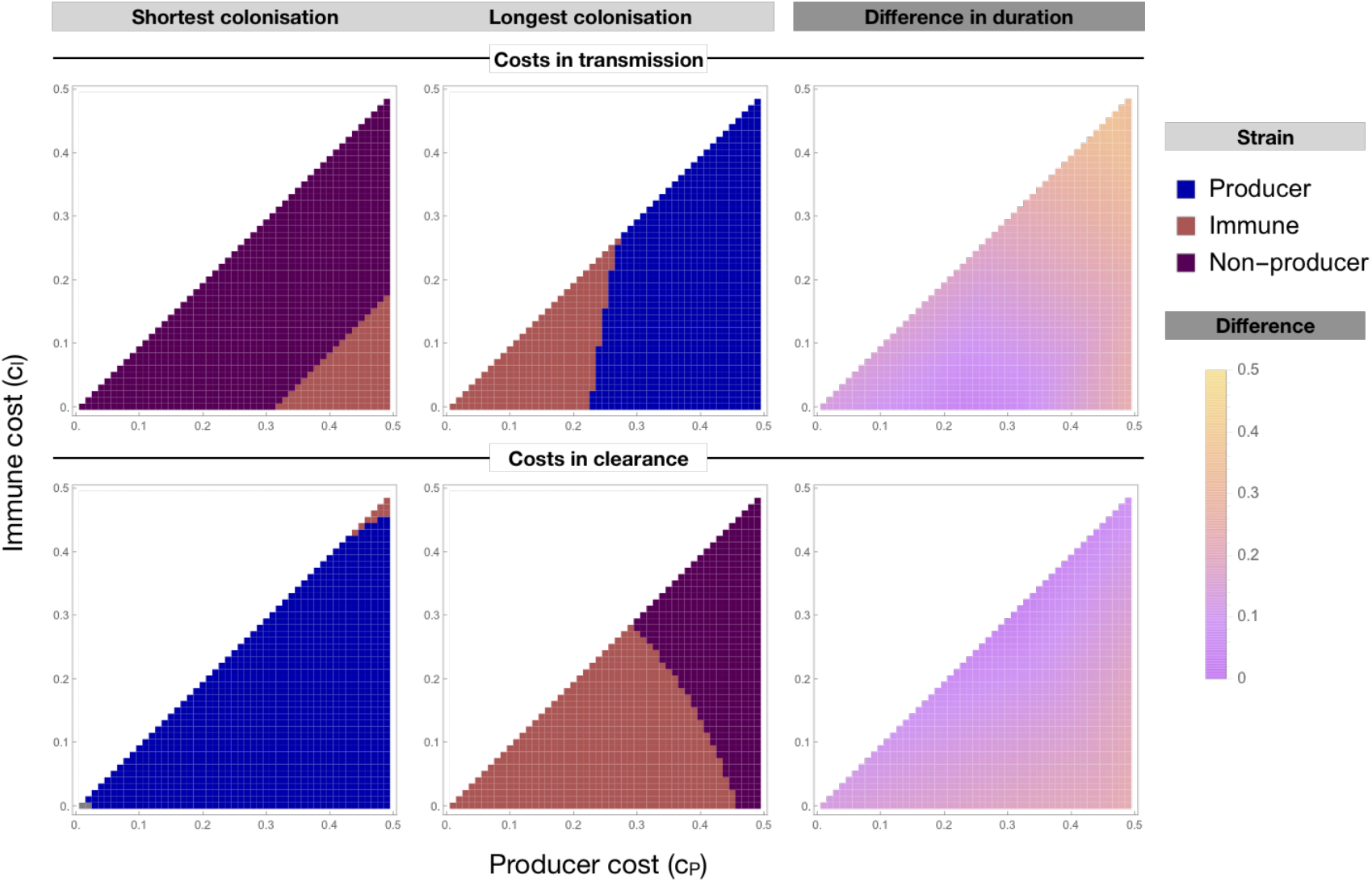
Variation in the duration of colonisation (i.e. reciprocal of the overall clearance rate) of strains with different bacteriocin profiles as a function of the costs associated with immunity and bacteriocin production. For each panel, the x-axis represents the fitness cost incurred by the producer strain; the y-axis the fitness cost incurred by the immune strain. The left and middle panels show which strain is associated with the longest and shortest duration of colonisation, respectively. The right panel shows the range of duration of colonisation (i.e. difference between the longest and shortest duration of colonisation). In the top panels, fitness costs decrease transmission, in the bottom panels, costs increase clearance rate. Other parameter values are *β* = 3, *µ* = 1, *k*_*MP*_ = *k*_*NM*_ = 0.5, *k*_*P N*_ = 1 and all other *k* = 0. For results on the duration of colonisation associated with each strain as well as results for other parameter values, see SI Section 2.5.

#### The effect of bacteriocins on antibiotic resistance dynamics

The observed variation in the duration of colonisation of different bacteriocin profiles suggests that diversity in bacteriocin profiles could indeed maintain diversity at the antibiotic resistance locus. To investigate the effect of bacteriocins on antibiotic resistance frequencies in more detail, we expand the model of bacteriocin dynamics (Equations 1) to include competition between resistant and sensitive strains.

We model a species that is carried asymptomatically most of the time (e.g. *S. pneumoniae*)— the antibiotic exposure of hosts is therefore independent of whether they are colonised and equal to the average antibiotic consumption rate in the population (*τ*). Interactions between the three bacteriocin profiles are the same as previously described. In addition, each profile can be either antibiotic sensitive (*S*) or antibiotic resistant (*R*), giving rise to six possible strains. Antibiotic sensitive strains are subject to an additional clearance rate *τ* (we assume immediate clearance in response to antibiotics). Resistance carries a fitness cost, which can be associated with clearance and/or transmission rate. The full model structure is given in SI Section 1.2.

Consistent with previous results, the between-strain variation in duration of colonisation allows coexistence of antibiotic sensitivity and resistance. Antibiotic resistance is associated with the bacteriocin profile(s) with longer duration of colonisation (Figure 5). The range of antibiotic consumption rates over which coexistence is observed and the frequency of antibiotic resistance in this region of coexistence is highly dependent on parameters. The finding that diversity at the bacteriocin locus can maintain intermediate antibiotic resistance frequencies is generally robust, with the exception of some specific cases when the cost of antibiotic resistance affects clearance rate. These are discussed in detail in SI Section 2.6.

**Figure 5:**
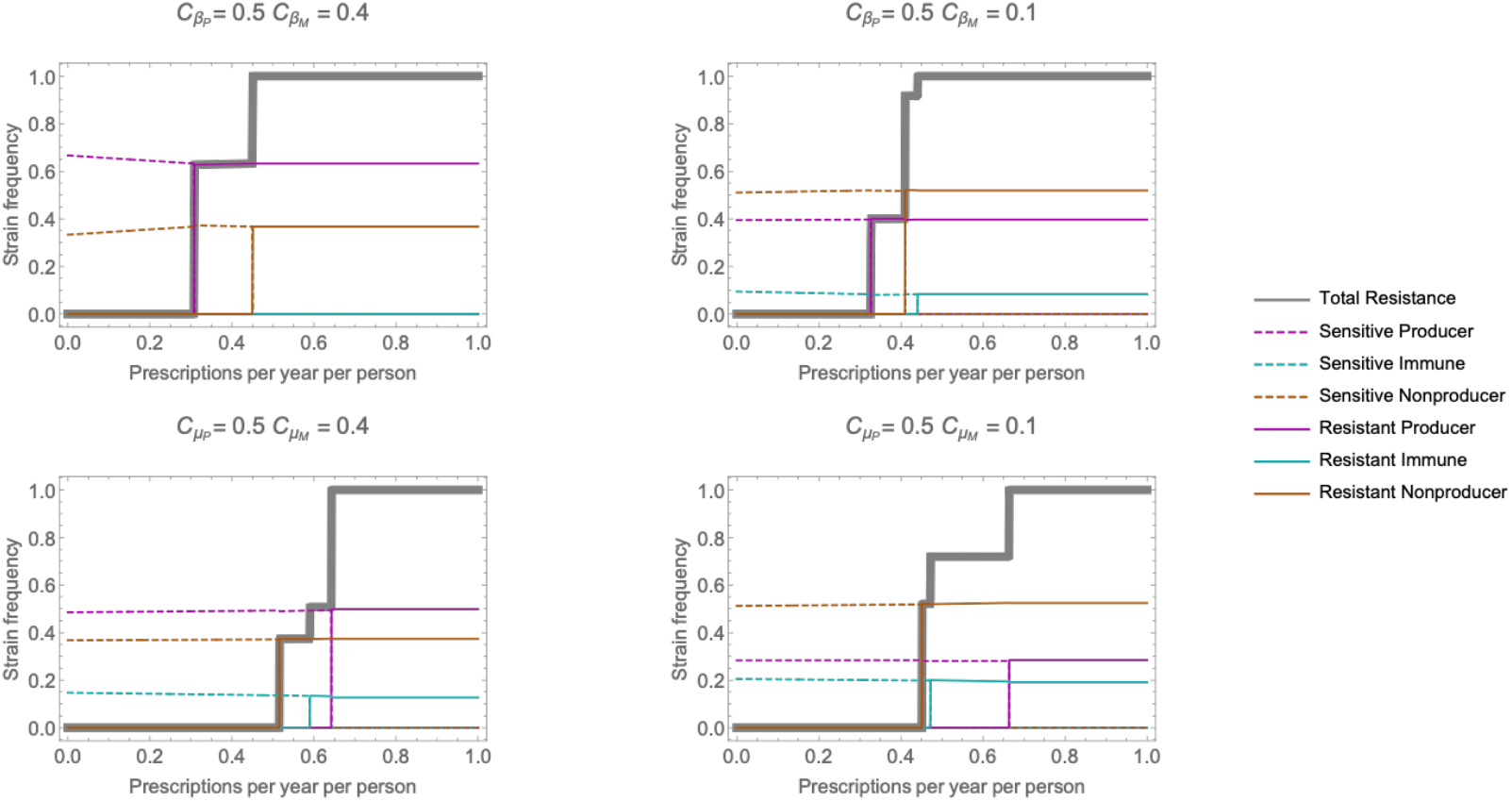
Strain frequencies (colours) and overall resistance frequency (gray) as a function of the population antibiotic consumption rate (*τ*) for different costs of the producer and immune strains. *c*_*β*_ indicates cost in transmission and *c*_*µ*_ indicates cost in clearance, with *P* and *M* indicating the cost to the producer and immune strain respectively: in the top row, bacteriocin-associated costs affect transmission, in the bottom row these affect clearance. The cost of antibiotic resistance is modelled as affecting transmission (see SI Section 2.6 for antibiotic resistance costs modelled as affecting clearance). Other parameters are *β* = 3, *µ* = 1, *k*_*P N*_ = 1, *k*_*MP*_ = *k*_*NM*_ = 0.5. Due to the difficulty of computing equilibrium solutions and eigenvalues for this model, these results are derived through simulating rather than stability analysis (simulation until *t* = 10^5^ months).

## Discussion

This paper was motivated by two main questions: firstly, the role of epidemiological processes in bacteriocin dynamics and diversity in *S. pneumoniae* and similar species; and secondly, whether these epidemiological processes could generate variation in duration of colonisation between different bacteriocin profiles and, as a result, contribute to the observed coexistence of antibiotic sensitive and resistant strains.

We have shown that bacteriocin diversity is readily maintained in epidemiological models. This finding is robust to assumptions about how the inhibitory activity of bacteriocins and costs associated with bacteriocin production and immunity translate to the epidemiological scale (e.g. within-host fitness differences leading to one strain displacing another vs differences in epidemiological parameters vs a mixture of the two): coexistence of different bacteriocin profiles is found under a range of assumptions. We find that the specifics of this coexistence (i.e. which bacteriocin profiles coexist and at what frequencies) are highly dependent on parameter values. This sensitivity to the properties of the bacteriocins—such as the cost of toxin production—may explain the observed variability in characteristics (e.g. prevalence and composition of bacteriocin loci) of bacteriocin systems.

In addition to the robust diversity in bacteriocin profiles in epidemiological models, we can also draw other general insights from these results.

Firstly, the inclusion of transmission dynamics introduces additional mechanisms through which strain diversity can be maintained, allowing different combinations of strains to coexist. Previous work had suggested that all three strains (producer, non-producer and immune) must be present to allow diversity to persist: the maintenance of diversity depended on a non-transitive competitive structure (‘rock-paper-scissors dynamics’) [14, 17]. In our epidemiological model, we also observe coexistence of the producer and non-producer strain without the immune strain. The mechanism at play here is not rock-paper-scissors dynamics but rather a competition-colonisation trade-off [30]: the producer strain has an advantage at the within-host scale (i.e. it can displace the non-producer) but a disadvantage at the between-host scale (i.e. it pays a fitness cost which reduces transmission rate and/or increases clearance rate). This type of trade-off gives rise to negative frequency-dependent selection—the benefit of the within-host advantage depends on the prevalence of the non-producer strain—and thus maintains coexistence of the competitors [31].

Secondly, in *S. pneumoniae*, some bacteriocin systems, such as the *pld* locus [11] and the circular toxin pneumocyclicin [12], are found on only a subset of strains. Others, such as the *blp* [4, 5, 6, 7] and *cib* [8] loci, are ubiquitous. Such ubiquity is not consistent with our epidemiological model under standard assumptions about bacteriocin ecology: all stable equilibria require the presence of the non-producer strain, because toxin production is only beneficial when susceptible strains are present. Our results highlight two mechanisms which allow toxin-producer to exist in absence of non-producers (in other words, mechanisms that allow the bacteriocin locus to fully invade the bacterial population). Firstly, exclusion of the non-producer strain is seen when the toxins provide an advantage in both invasion of already colonised hosts *and* colonisation of uncolonised hosts (a biological interpretation would be that toxins are effective against both the focal *and* other species). Toxin production is maintained in absence of the non-producer strain because it provides a benefit despite the absence of the non-producer. Secondly, in the pherotype model, immunity is not protective against toxins released by a different pherotype, and strains of one pherotype are therefore susceptible to toxins from another pherotype. Toxin production is therefore beneficial even in absence of the non-producer strain. Indeed, the ubiquitious *cib* and *blp* loci are both associated with a signalling locus (i.e. pherotype), suggesting this may be the explanation for their ubiquity.

In the second part of the paper, we have shown that bacteriocins are a plausible candidate for involvement in resistance dynamics. We predict differences in duration of colonisation of different bacteriocin profiles, with the magnitude of these differences is highly sensitive to parameter values. The duration of colonisation is predicted to modulate the fitness of antibiotic resistance [24]; we therefore expect an association between antibiotic resistance and bacteriocin profile and indeed observe this in a model incorporating both bacteriocin and resistance dynamics.

Testing this predicted association empirically is possible in theory, but would prove challenging given our current understanding of bacteriocin systems. Firstly, the direction of association between bacteriocin profile and antibiotic resistance depends on parameters. It is therefore unclear whether we expect antibiotic resistance to be associated with the producer, non-producer or immune phenotype, and indeed, this may differ between different bacteriocin systems. Secondly, the mapping between these modelled phenotypes and observed genotypes is non-trivial: although toxin and immunity genes can be identified, the effect on phenotype is less clear when multiple toxin and immunity genes are present on a genome (either because of the presence of a multi-toxin bacteriocin system or the presence of multiple different systems on one genome). Furthermore, in some bacteriocin systems, additional factors may affect the association between toxin and immunity genes; for example, a large proportion of *blp* systems may not be able to secrete pheromones and toxins because of an impaired transporter [32] and there is evidence of regulatory interplay between the *blp* and *cib* systems [33, 34]. Thus, although we have shown bacteriocins are a plausible candidate for involvement in resistance dynamics, more specific predictions about this involvement will require a more detailed understanding of specific bacteriocin systems.

It is worth highlighting two potential limitations of our modelling approach. Firstly, we do not model co-colonisation: within-host strain displacement is assumed to be fast. This assumption may not hold: co-colonisation with multiple strains is known to occur (e.g. 25% to 50% of colonised hosts depending on setting [19, 20, 21]), although whether such co-colonisation is possible between strains with different bacteriocin profiles is not clear. Furthermore, previous theoretical work suggests that assumptions about within-host dynamics and competition can have considerable impact on predictions about strain diversity at the between-host scale [35, 36, 37]. We tested the impact of allowing slower within-host dynamics and thus co-colonisation in the simplest version of the model—where strain displacement is only possible through bacteriocin action (Figure 2 B)—and find our results are indeed qualitatively robust (SI Section 2.1). However, it remains possible that the impact of co-colonisation is more significant in more complex versions of the model, in particular the multi-locus extensions (i.e. pherotype and antibiotic resistance models).

Secondly, we consider toxin-production and immunity (and antibiotic resistance) as binary traits: strains either possesses genes encoding for toxins and immunity or they do not. This modelling approach reflects observed variation in the absence/presence of toxin and immunity genes in pneumococcal genomes [4, 5, 6, 7, 8, 11, 12]. An alternative approach would be to treat these traits as continuous variables; this would correspond to assuming genes are present on all strains and that effects are tunable (e.g. through modulation of gene expression), leading to different levels of toxicity, immunity and fitness cost. This would allow modelling the evolution of the level of toxicity and immunity. Such approach would require assumptions about how tunable bacteriocin effects are; trade-offs between fitness cost and levels of toxicity and immunity; and the relative time-scale of evolutionary and epidemiological processes. Such assumptions will be subject to considerable uncertainty. For example, the extent to which bacteriocins are tunable is unclear—indeed, the relationship between inhibition and toxin concentration appears threshold-like rather than gradual [7], suggesting inhibitory effects may not be readily modulated. Nevertheless, the impact of assuming bacteriocin traits are evolvable on a similar time scale to epidemiological processes is an open and interesting question.

In summary, we have shown that diversity in bacteriocin profiles arises robustly in epidemiological models of bacteriocin dynamics and can be maintained through either rock-paper-scissors dynamics or a colonisation-competition trade-off. The specifics of the predicted diversity are sensitive to assumptions about how bacteriocins affect epidemiological processes and on bacteriocin-related parameters (e.g. costs of bacteriocin production and immunity), providing a potential explanation for differences between bacteriocin systems. We have also demonstrated that diversity at bacteriocin loci is a plausible candidate for also maintaining diversity at resistance loci. These insights arise from modelling that approaches bacteriocins dynamics from a high level of abstraction, rather than representing a specific bacteriocin system. Generating more specific insights into particular bacteriocins systems will require models informed by the biology of the specific system. Therefore, a more complete understanding of the role of bacteriocins in bacterial ecology will need a more specific characterisation of their effects on transmission, invasion, within-host competition and duration of colonisation.

## Acknowledgments

We thank Marc Lipsitch for comments on an earlier version of the manuscript. This work was funded by the National Institutes of Health (MIDAS) (SL and CF grant ref: U01GM110721-01), the Li Ka Shing foundation (SL and CF); and a Sir Henry Dale Fellowship, jointly funded by Wellcome and the Royal Society (NJC, grant ref: 104169/Z/14/Z)

## References

[1] Hawlena H, Bashey F, Lively CM. Bacteriocin-mediated interactions within and between coexisting species. Ecology and Evolution. 2012;2(10):2521–2526.

[2] Riley MA, Wertz JE. Bacteriocins: evolution, ecology, and application. Annual Reviews in Microbiology. 2002;56(1):117–137.

[3] Miller EL, Abrudan MI, Roberts IS, Rozen DE. Diverse ecological strategies are encoded by Streptococcus pneumoniae bacteriocin-like peptides. Genome biology and evolution. 2016;8(4):1072–1090.

[4] de Saizieu A, Gardès C, Flint N, Wagner C, Kamber M, Mitchell TJ, et al. Microarray-based identification of a novelStreptococcus pneumoniae regulon controlled by an autoinduced peptide. Journal of Bacteriology. 2000;182(17):4696–4703.

[5] Reichmann P, Hakenbeck R. Allelic variation in a peptide-inducible two-component system of Streptococcus pneumoniae. FEMS microbiology letters. 2000;190(2):231–236.

[6] Dawid S, Roche AM, Weiser JN. The blp bacteriocins of Streptococcus pneumoniae mediate intraspecies competition both in vitro and in vivo. Infection and Immunity. 2007;75(1):443–451.

[7] Lux T, Nuhn M, Hakenbeck R, Reichmann P. Diversity of Bacteriocins and Activity Spectrum in Streptococcus pneumoniae. Journal of Bacteriology. s2007 Nov;189(21):7741–7751.

[8] Guiral S, Mitchell TJ, Martin B, Claverys JP. Competence-programmed predation of noncompetent cells in the human pathogen Streptococcus pneumoniae: genetic require-ments. Proceedings of the National Academy of Sciences of the United States of America. 2005;102(24):8710–8715.

[9] Croucher NJ, Coupland PG, Stevenson AE, Callendrello A, Bentley SD, Hanage WP. Diver-sification of bacterial genome content through distinct mechanisms over different timescales. Nature communications. 2014;5(1):1–12.

[10] Miller EL, Kjos M, Abrudan MI, Roberts IS, Veening JW, Rozen DE. Eavesdropping and crosstalk between secreted quorum sensing peptide signals that regulate bacteriocin production in Streptococcus pneumoniae. The ISME journal. 2018;12(10):2363–2375.

[11] Maricic N, Anderson ES, Opipari AE, Emily AY, Dawid S. Characterization of a Multi-peptide Lantibiotic Locus in Streptococcus pneumoniae. mBio. 2016;7(1):e01656–15.

[12] Bogaardt C, van Tonder AJ, Brueggemann AB. Genomic analyses of pneumococci reveal a wide diversity of bacteriocins–including pneumocyclicin, a novel circular bacteriocin. BMC Genomics. 2015;16(1):554.

[13] Levin B. Frequency-dependent selection in bacterial populations. Philosophical Transactions of the Royal Society of London B, Biological Sciences. 1988;319(1196):459–472.

[14] Durrett R, Levin S. Allelopathy in spatially distributed populations. Journal of Theoretical Biology. 1997;185(2):165–171.

[15] Frean M, Abraham ER. Rock–scissors–paper and the survival of the weakest. Proceedings of the Royal Society of London Series B: Biological Sciences. 2001;268(1474):1323–1327.

[16] Czárán TL, Hoekstra RF, Pagie L. Chemical warfare between microbes promotes biodiversity. Proceedings of the National Academy of Sciences. 2002;99(2):786–790.

[17] Kerr B, Riley MA, Feldman MW, Bohannan BJ. Local dispersal promotes biodiversity in a real-life game of rock–paper–scissors. Nature. 2002;418(6894):171.

[18] Kelsic ED, Zhao J, Vetsigian K, Kishony R. Counteraction of antibiotic production and degradation stabilizes microbial communities. Nature. 2015;521(7553):516.

[19] Hjálmarsdóttir MÁ, Gumundsdóttir PF, Erlendsdóttir H, Kristinsson KG, Haraldsson G. Cocolonization of pneumococcal serotypes in healthy children attending day care centers: molecular versus conventional methods. The Pediatric infectious disease journal. 2016;35(5):477–480.

[20] Kamng’ona AW, Hinds J, Bar-Zeev N, Gould KA, Chaguza C, Msefula C, et al. High multiple carriage and emergence of Streptococcus pneumoniae vaccine serotype variants in Malawian children. BMC Infectious Diseases. 2015;15(1):234.

[21] Turner P, Hinds J, Turner C, Jankhot A, Gould K, Bentley SD, et al. Improved detection of nasopharyngeal cocolonization by multiple pneumococcal serotypes by use of latex agglutination or molecular serotyping by microarray. Journal of Clinical Microbiology. 2011;49(5):1784–1789.

[22] Shen P, Lees JA, Bee GCW, Brown SP, Weiser JN. Pneumococcal quorum sensing drives an asymmetric owner–intruder competitive strategy during carriage via the competence regulon. Nature Microbiology. 2019;4(1):198–208.

[23] Valente C, Dawid S, Pinto FR, Hinds J, Simões AS, Gould KA, et al. The blp locus of Streptococcus pneumoniae plays a limited role in the selection of strains that can cocolonize the human nasopharynx. Appl Environ Microbiol. 2016;82(17):5206–5215.

[24] Lehtinen S, Blanquart F, Croucher NJ, Turner P, Lipsitch M, Fraser C. Evolution of antibiotic resistance is linked to any genetic mechanism affecting bacterial duration of carriage. Proceedings of the National Academy of Sciences of the United States of America. 2017 jan;114(5):1075–1080.

[25] Lehtinen S, Blanquart F, Lipsitch M, Fraser C, with the Maela Pneumococcal Collaboration, et al. On the evolutionary ecology of multidrug resistance in bacteria. PLoS Pathogens. 2019;15(5):e1007763.

[26] Colijn C, Cohen T, Fraser C, Hanage W, Goldstein E, Givon-Lavi N, et al. What is the mechanism for persistent coexistence of drug-susceptible and drug-resistant strains of Streptococcus pneumoniae? Journal of The Royal Society Interface. 2010;7(47).

[27] Lees JA, Croucher NJ, Goldblatt D, Nosten F, Parkhill J, Turner C, et al. Genome-wide identification of lineage and locus specific variation associated with pneumococcal carriage duration. Elife. 2017;6:e26255.

[28] Inc WR. Mathematica, Version 11.2;. Champaign, IL, 2017.

[29] Lehtinen S, Chewapreecha C, Lees J, Hanage WP, Lipsitch M, Croucher NJ, et al. Horizontal gene transfer rate is not the primary determinant of observed antibiotic resistance frequencies in Streptococcus pneumoniae. Science advances. 2020;6(21):eaaz6137.

[30] Leibold MA, Holyoak M, Mouquet N, Amarasekare P, Chase JM, Hoopes MF, et al. The metacommunity concept: a framework for multi-scale community ecology. Ecology Letters. 2004;7(7):601–613.

[31] Alizon S. Co-infection and super-infection models in evolutionary epidemiology. Interface focus. 2013;3(6):20130031.

[32] Son MR, Shchepetov M, Adrian PV, Madhi SA, de Gouveia L, von Gottberg A, et al. Conserved mutations in the pneumococcal bacteriocin transporter gene, blpA, result in a complex population consisting of producers and cheaters. MBio. 2011;2(5):e00179–11.

[33] Wholey WY, Kochan TJ, Storck DN, Dawid S. Coordinated bacteriocin expression and competence in Streptococcus pneumoniae contributes to genetic adaptation through neighbor predation. PLoS pathogens. 2016;12(2):e1005413.

[34] Kjos M, Miller E, Slager J, Lake FB, Gericke O, Roberts IS, et al. Expression of Streptococcus pneumoniae bacteriocins is induced by antibiotics via regulatory interplay with the competence system. PLoS pathogens. 2016;12(2):e1005422.

[35] Davies NG, Flasche S, Jit M, Atkins KE. Within-host dynamics shape antibiotic resistance in commensal bacteria. Nature Ecology & Evolution. 2019;3(3):440.

[36] Mulberry N, Rutherford A, Colijn C. Systematic comparison of coexistence in models of drug-sensitive and drug-resistant pathogen strains. Theoretical Population Biology. 2019;.

[37] Igler C, Huisman JS, Siedentop B, Bonhoeffer S, Lehtinen S. Plasmid co-infection: linking biological mechanisms to ecological and evolutionary dynamics. Philosophical Transactions of the Royal Society B. 2022;377(1842):20200478.

