## Supporting Information for "Epidemiological dynamics of bacteriocin competition and antibiotic resistance"

#### 1 Additional models

##### 1.1 Pherotype model

In the pherotype model, we consider the interactions between strains of pherotype A and B. Each bacteriocin profile can have either pherotype, giving rise to a model with six strains: $I_{P^A}$, $I_{M^A}$, $I_{N^A}$, $I_{P^B}$, $I_{M^B}$, $I_{N^B}$. For simplicity, we assume bacteriocins are required for invasion (i.e. similar to the model structure in main text Figure 2 B). Within pherotype, the interactions remain as described in the main text. Immunity is not effective across pherotypes, allowing a producer strain to invade an immune strain of a different pherotype. We also assume that the $k$ parameter is the same for all invasions ($k = k_{P^A N^A} = k_{P^A N^B} = k_{P^A M^B} = k_{P^B N^B} = k_{P^B N^A} = k_{P^B M^A}$). This results in the following dynamics, with $X = 1 - I_{P^A} - I_{M^A} - I_{N^A} - I_{P^B} - I_{M^B} - I_{N^B}$:

$$\begin{aligned}
\frac{dI_{P^A}}{dt} &= \beta_P I_{P^A} X + k\beta_P (I_{N^A} + I_{N^B} + I_{M^B}) I_{P^A} - \mu_P I_{P^A} \\
\frac{dI_{M^A}}{dt} &= \beta_M I_{M^A} X - k\beta_P (I_{P^B}) - \mu_M I_{M^A} \\
\frac{dI_{N^A}}{dt} &= \beta_N I_{N^A} X - k\beta_P (I_{P^A} + I_{P^B}) - \mu_N I_{N^A} \\
\frac{dI_{P^B}}{dt} &= \beta_P I_{P^B} X + k\beta_P (I_{N^A} + I_{N^B} + I_{M^A}) I_{P^B} - \mu_P I_{P^B} \\
\frac{dI_{M^B}}{dt} &= \beta_M I_{M^B} X - k\beta_P (I_{P^A}) - \mu_M I_{M^B} \\
\frac{dI_{N^B}}{dt} &= \beta_N I_{N^B} X - k\beta_P (I_{P^A} + I_{P^B}) - \mu_N I_{N^B}
\end{aligned} \tag{1}$$

There are two things to note about how this model formulation. Firstly, whether a producer strain is able to invade a host colonised with a producer strain of the other pherotype would dependent on assumptions about the timing of toxin and immunity expression. However, this does not impact model dynamics: the rate at which the pherotype A producer replaces the pherotype B producer is equal to the rate at which the pherotype B producer replaces the pherotype A producer. These terms therefore cancel out and we have not included them in SI Equations 1.

Secondly, the non-producer strain is always susceptible to the toxin—producer strains can therefore replace non-producer strains regardless of pherotype. Non-producers of the two pherotypes are therefore ecologically indistinguishable and modelling the two non-producers strains as a single strain would give identical results.

##### 1.2 Resistance model

We model a species that is carried asymptomatically most of the time (e.g. *S. pneumoniae*)—the antibiotic exposure of hosts is therefore independent of whether they are colonised and simply equal to antibiotic consumption rate in the population ($\tau$). Interactions between the three bacteriocin profiles are the same as in the main text model. In addition, each profile can be either antibiotic sensitive ($S$) or antibiotic resistant ($R$), giving rise to six possible strains ($I_{P^S}$, $I_{M^S}$, $I_{N^S}$, $I_{P^R}$, $I_{M^R}$, $I_{N^R}$). Antibiotic sensitive strains are subject to an additional clearance rate $\tau$ (we assume immediate clearance in response to antibiotics). Resistance carries a fitness cost ($c_R$), which can affect either transmission and/or clearance. Resistance- and bacteriocin-associated costs are modelled as multiplicative: e.g. $\beta_{P^R} = (1 - c_P)(1 - c_R)\beta$ , where $\beta$ is the baseline transmission rate (i.e. the transmission rate of the sensitive non-producer strain $\beta_{N^S}$).

The dynamics of this system are described by SI Equations 2. For clarity, we have set $k_{PM} = k_{MN} = k_{NP} = 0$: strain displacement is only possible when the invading strain is fitter (as is the case for all of the results presented in the main text). With $X = 1 - I_{P^S} - I_{M^S} - I_{N^S} - I_{P^R} - I_{M^R} - I_{N^R}$, this gives:

$$\begin{aligned}
\frac{dI_{P^S}}{dt} &= \beta_{P^S} I_{P^S} X - k_{MP}(\beta_{M^S} I_{M^S} + \beta_{M^R} I_{M^R}) I_{P^S} + k_{PN}\beta_{P^S}(I_{N^S} + I_{N^R}) I_{P^S} - (\mu_{P^S} + \tau) I_{P^S} \\
\frac{dI_{M^S}}{dt} &= \beta_{M^S} I_{M^S} X - k_{NM}(\beta_{N^S} I_{N^S} + \beta_{N^R} I_{N^R}) I_{M^S} + k_{MP}\beta_{M^S}(I_{P^S} + I_{P^R}) I_{M^S} - (\mu_{M^S} + \tau) I_{M^S} \\
\frac{dI_{N^S}}{dt} &= \beta_{N^S} I_{N^S} X - k_{PN}(\beta_{P^S} I_{P^S} + \beta_{P^R} I_{P^R}) I_{N^S} + k_{NM}\beta_{N^S}(I_{M^S} + I_{M^R}) I_{N^S} - (\mu_{N^S} + \tau) I_{N^S} \\
\frac{dI_{P^R}}{dt} &= \beta_{P^R} I_{P^R} X - k_{MP}(\beta_{M^S} I_{M^S} + \beta_{M^R} I_{M^R}) I_{P^R} + k_{PN}\beta_{P^R}(I_{N^S} + I_{N^R}) I_{P^R} - \mu_{P^R} I_{P^R} \\
\frac{dI_{M^R}}{dt} &= \beta_{M^R} I_{M^R} X - k_{NM}(\beta_{N^S} I_{N^S} + \beta_{N^R} I_{N^R}) I_{M^R} + k_{MP}\beta_{M^R}(I_{P^S} + I_{P^R}) I_{M^R} - \mu_{M^R} I_{M^R} \\
\frac{dI_{N^R}}{dt} &= \beta_{N^R} I_{N^R} X - k_{PN}(\beta_{P^S} I_{P^S} + \beta_{P^R} I_{P^R}) I_{N^R} + k_{NM}\beta_{N^R}(I_{M^S} + I_{M^R}) I_{N^R} - \mu_{N^R} I_{N^R}
\end{aligned} \tag{2}$$

### 2 Additional text and results

#### 2.1 Slow within-host dynamics

In the main text, we make the assumption that bacteriocin interactions within-host lead to rapid strain replacement within-the host: we do not model co-colonised states. This assumption may not hold: co-colonisation with multiple strains is known to occur (e.g. 25% to 50% of colonised hosts depending on setting [1, 2, 3]), although whether such co-colonisation can occur between strains with different bacteriocin profiles is not clear. To verify the robustness of our results to relaxing this assumption about rapid within-host dynamics, we expand our model to include a co-colonised compartment. For simplicity, we assume invasion only occurs between producer and non-producers strains ($k_{PN} \geq 0$, all other $k$ are 0, as in main text Figure 2 B). The model is identical to the main text, apart from the addition of category of hosts co-colonised with $P$ and $N$. Hosts colonised with $N$ become co-colonised when invaded with $P$ (at rate $k_{PN}\beta_P P$). Co-colonised host become singly colonised with $P$ as the producer out-competes the non-producer (at rate $r$). We assume co-colonised hosts transmit each strain at half the rate of an individual singly colonised with the strain. This gives rise to the following model, with $X = 1 - I_P - I_M - I_N - I_{PN}$:

$$\begin{aligned}
\frac{dI_P}{dt} &= \beta_P(I_P + I_{PN}/2)X + rI_{PN} - \mu_P I_P \\
\frac{dI_M}{dt} &= \beta_M I_M X - \mu_M I_M \\
\frac{dI_N}{dt} &= \beta_N(I_N + I_{PN}/2)X - k_{PN}\beta_P(I_P + I_{PN}/2)I_N - \mu_N I_N \\
\frac{dI_{PN}}{dt} &= k_{PN}\beta_P(I_P + I_{PN}/2)I_N - rI_{PN} - \mu_{PN} I_{PN}
\end{aligned} \tag{3}$$

To explore the impact of slow dynamics, we vary the speed of the within-host dynamics (i.e. the $r$ parameters, with larger $r$ corresponding to faster dynamics) and compare the results to the main text model. Strain coexistence remains qualitatively similar to the outcomes of the main text model (SI Figure S1). Slower within-host dynamics benefit the non-producer (as the producer is less effective at out-competing it) and thus lead to a larger parameter space in which the non-producer strain excludes the other two strains.

#### 2.2 Defensive bacteriocins

In the main text, bacteriocins are generally conceptualised as offensive: fitness differences between strains (i.e. $P$ out-competing $N$, $N$ out-competing $M$, $M$ out-competing $P$) allow the

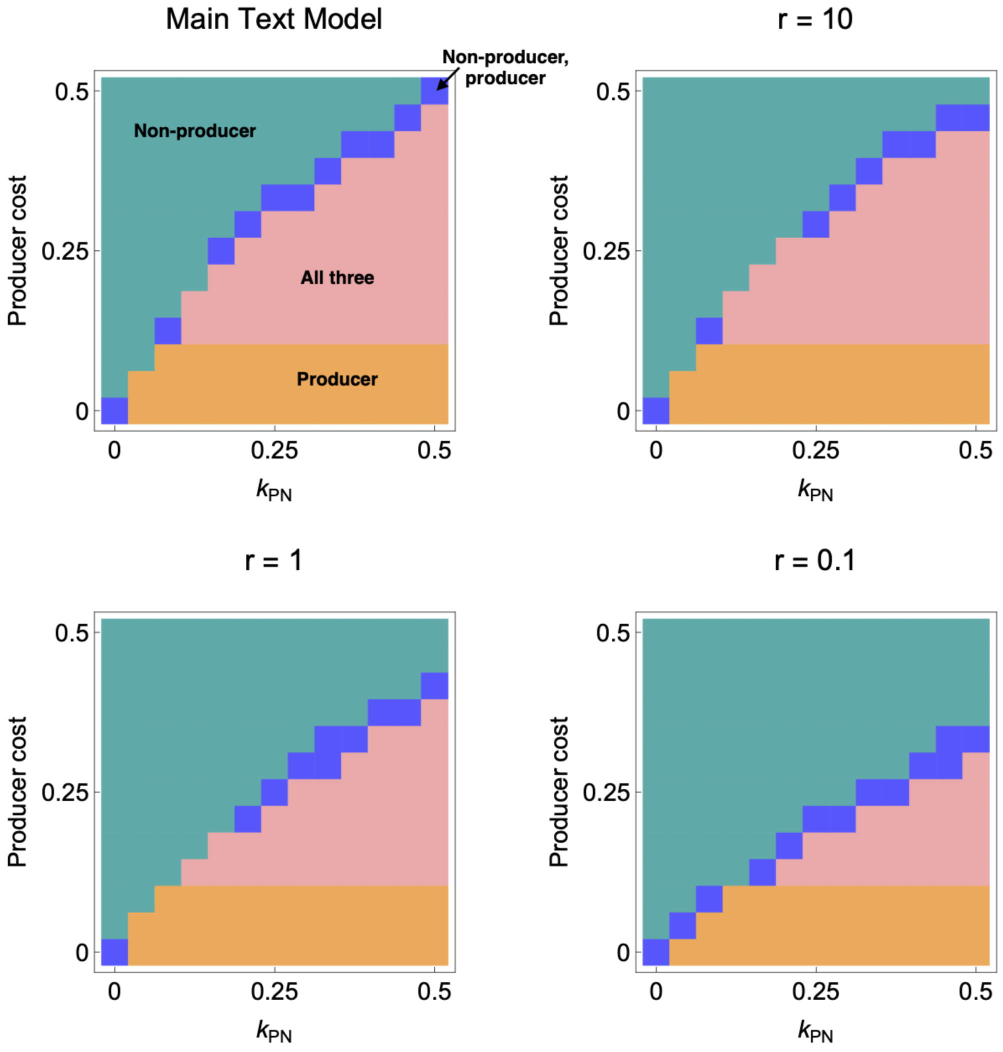


Figure S1: Qualitative outcomes are robust to relaxing assumption about rapid within-host dynamics: comparison of the main text model with models with an additional category of hosts co-colonised with the producer and non-producer. The $r$ parameter captures the rate at which $P$ out-competes $N$ within co-colonised hosts. The x-axis captures the rate at which the producer co-colonises the non-producer. The y-axis captures the transmission cost ($c_P$) to the producer strain ($\beta_P = (1 - c_P)\beta$). The colours indicate which strains are present at equilibrium, with green corresponding to the non-producer only, blue to the non-producer and producer, pink to all three and orange to the producer only. Unlike the main text results, the strains present at equilibrium are determined by simulating the model until $t = 10^5$ months, as computing the eigenvalues for this modified model was prohibitively computationally expensive. Parameters are $\beta = \beta_N = \beta_M = 3$, $\mu = 1$ and $k = 0$ for all $k$ other than $k_{PN}$.

fitter strain to invade a host colonised by the less fit stain. Bacteriocins could also be conceptualised as defensive—in this case, fitness differences would allow a fitter strain to resist invasion by a less fit strain—or a combination of the two. As discussed in the main text, when strains have identical transmission rates, these views are mathematically equivalent. If transmission rates are not equal due to fitness costs associated with transmission, modelling bacteriocins as defensive is not equivalent to the offensive model of the main text. This is because the rate of successful invasion by strain $A$ depends on $\beta_A$, whereas the rate of strain $A$ being invaded by strain $B$ depends on $\beta_B$. As a result, the interaction between strains $A$ and $B$ in the dynamics of strain $A$ $[(\beta_A k_{AB} - \beta_B k_{BA}) I_A I_B]$ does not cancel out. SI Figure S2 shows that results relating to strain diversity are qualitatively similar to those presented in the main text when transmission rates are not equal and bacteriocins are modelled as defensive instead of offensive.

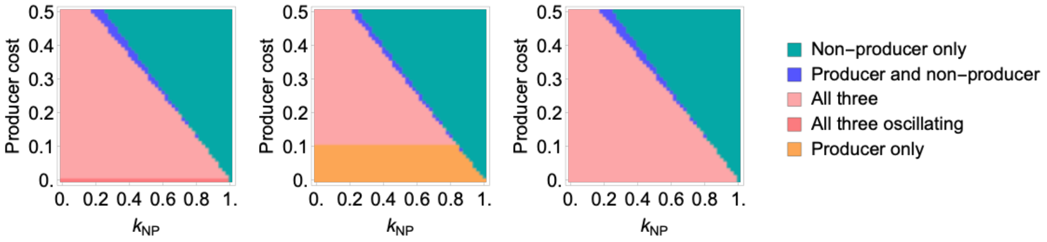


Figure S2: Qualitative outcomes are similar when bacteriocins are modelled as defensive rather than offensive. For all plots, the x-axis captures how well the producer strain resists invasion from the non-producer and the y-axis captures the transmission cost to the producer strain ($\beta_P = (1 - c_P)\beta_N$). For all plots, $\mu = 1$, $\beta_N = 3$, and $k_{MP} = k_{PN} = k_{NM} = 1$ **Left** $k_{MN} = k_{PM} = 0.5$, $\beta_M = 3$ **Middle** $k_{MN} = k_{PM} = 1$, $\beta_M = 2.7$ **Right** $k_{MN} = k_{PM} = 0.5$, $\beta_M = 2.7$.

#### 2.3 Differences in the properties of the two and three strain equilibria

Under standard assumptions about bacteriocins ecology (i.e. $P$ out-competes $N$, $N$ out-competes $M$ and $M$ out-competes $P$), the model admits two types of coexistence: either all three strains are present, or the producer and non-producer coexist without the immune strain. These two modes of coexistence arise from different mechanisms (see main text discussion): non-transitive rock-paper-scissors dynamics and a competition-colonisation trade-off. As a result, these equilibria behave differently: the three-strain equilibrium exhibits 'survival of the weakest' behaviour, where increasing the transmission rate or invasiveness of the producer strain *decreases* its equilibrium frequency (SI Figure S3). Note that this effect is specific to increasing the parameters which determine the rate at which the producer displaces the non-producer: increasing the fitness of the producer by decreasing its clearance rate has no effect on the producer's equilibrium frequency. Survival of the weakest behaviour is not observed in the two-strain equilibrium.

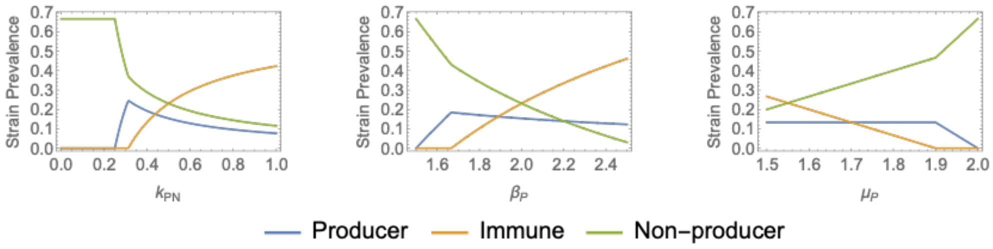


Figure S3: 'Survival of the weakest' behaviour is observed when all three strains are present. The plots show strain frequencies at equilibrium as a function of the competitiveness of the producer strain ($k_PN$, $\beta_P$ and $\mu_P$). For the first two parameters, increasing the fitness of the producer strain leads to a *decrease* in its equilibrium frequency when all three strains are present, but not when only two strains are present. This behaviour is not observed for the clearance rate. Parameters for each plot are: **left** $\beta_P = 2$, $\beta_M = 2.6$, $\beta_N = 3$, $\mu_P = \mu_M = \mu_N = 1$, all $k$ apart from $k_{PN}$ equal to 0; **middle** $\beta_M = 2.6$, $\beta_N = 3$, $\mu_P = \mu_M = \mu_N = 1$, $k_{PN} = 0.5$, all other $k = 0$; **right** $\beta_P = \beta_M = \beta_N = 3$, $\mu_M = 1.2$, $\mu_N = 1$, $k_{PN} = 0.5$, all other $k = 0$.

#### 2.4 Bacteriocin-associated costs in clearance

In the main text, bacteriocin-associated costs are modelled as a reduction in transmission rate. SI Figure S4 shows results are qualitatively similar when bacteriocin-associated costs are modelled as an increased clearance rate rather than reduced transmission rate.

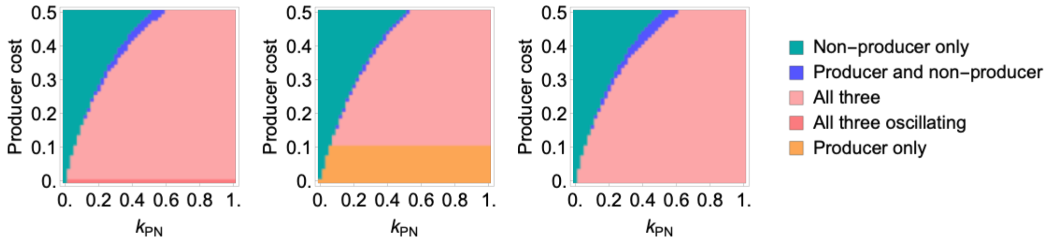


Figure S4: Qualitative outcomes are similar when bacteriocins costs are associated with clearance rather than transmission rate. For all plots, the x-axis captures how well the producer strain invades the non-producer and the y-axis captures the clearance cost to the producer strain ($\mu_P = \mu_N/(1 - c_P)$). For all plots, $\mu_N = 1$, $\beta_N = \beta_M = \beta_P = 3$, and $k_{PM} = k_{NP} = k_{MN} = 0$ **Left** $k_{MP} = k_{NM} = 0.5$, $\mu_M = 1$ **Middle** $k_{MP} = k_{MN} = 0$, $\mu_M = 10/9$ **Right** $k_{MP} = k_{NM} = 0.5$, $\mu_M = 10/9$.

#### 2.5 Variation in duration of colonisation

In the main text, we show which bacteriocin profiles have the longest and shortest duration of colonisation and the difference between the two durations. Figure S5 shows the same information but for different parameters than used in the main text, specifically lower invasion parameters ($k$). Figures S6 and S7 show the duration of colonisation associated with each strain for both parameter sets.

The durations of colonisation are sensitive to the costs associated with the producer and immune strain, as well as the invasion parameters ($k$). When costs affect transmission, variation in duration of colonisation arises solely from differences in the rate of strain displacement. The producer always has a longer duration of colonisation than the non-producer: the non-producer is more susceptible to displacement than the producer. For the explored parameter sets, the duration of colonisation of both the producer and non-producer increase with the producer cost. For the non-producer strain, the duration of colonisation increases because the rate at which the non-producer is displaced depends on the transmission rate of the producer and thus decreases with increasing transmission-associated costs. For the producer strain, the duration of colonisation increases because of the non-transitive competitive structure and 'survival of the weakest' effect: increasing the cost incurred by the producer decreases the frequency of the immune strain and thus decreases the rate at which the producer strain is replaced by the immune strain. Decreasing the invasion parameters increases duration of colonisation and thus decreases the extent of variation in duration of colonisation.

When costs are associated with clearance, variation in duration of colonisation arises both from variation in displacement and directly from the incurred costs. These two mechanisms affect the relative duration of colonisation of the producer and non-producer strains in opposite directions: strain displacement increases the colonisation duration of the producer relative to the non-producer, while the clearance-associated cost decreases the colonisation duration of the producer relative to the non-producer. For the explored parameter sets, the later has the greater effect and the producer therefore always has a shorter duration of colonisation than the non-producer. As a result of these opposing effects, when invasion parameters are large (Figure S6), the overall variation in duration of colonisation is smaller when costs are associated with clearance than when they are associated with transmission. Decreasing the invasion parameters increases the extent of variation in duration of colonisation (Figure S6).

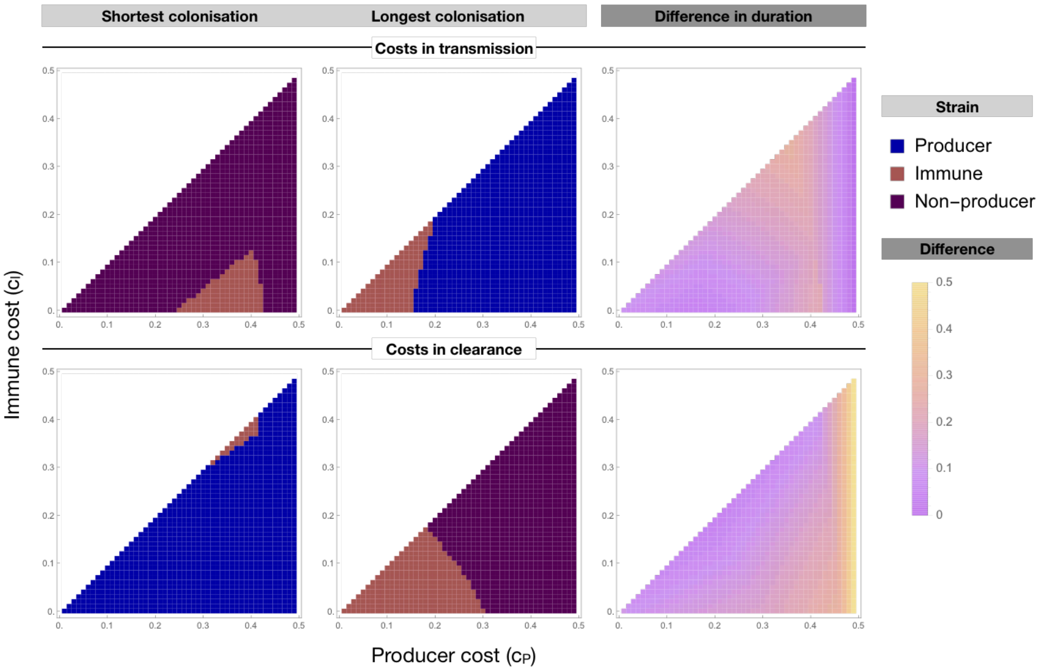


Figure S5: Variation in the duration of colonisation (i.e. reciprocal of the overall clearance rate) of strains with different bacteriocin profiles, for different invasion parameters than in main text Figure 4. As with the main text figure, for each panel, the x-axis represents the fitness cost incurred by the producer strain; the y-axis the fitness cost incurred by the immune strain. The left and middle panels show which strain is associated with the longest and shortest duration of colonisation, respectively. The right panel shows the range of duration of colonisation (i.e. difference between the longest and shortest duration of colonisation). In the top panels, fitness costs decrease transmission, in the bottom panels, costs increase clearance rate. Other parameter values are $\beta = 3$, $\mu = 1$, $k_{MP} = k_{NM} = 0.25$, $k_{PN} = 0.5$ and all other $k = 0$

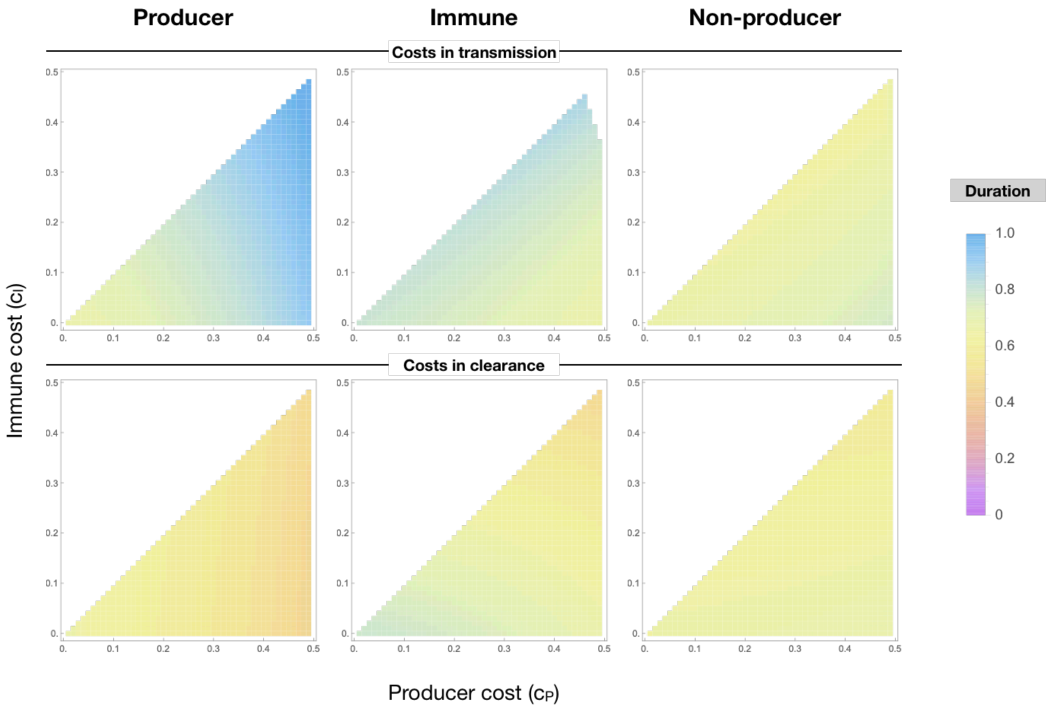


Figure S6: Duration of colonisation associated with each bacteriocin strain, for the same parameters as in main text Figure 4. As with the main text figure, for each panel, the x-axis represents the fitness cost incurred by the producer strain; the y-axis the fitness cost incurred by the immune strain. White space occurs when no duration is recorded. This happens when $c_M > c_P$ as we assume the producer strain incurs a higher cost than the immune strain, or when a particular strain is absent at equilibrium for particular values of fitness cost. In the top panels, fitness costs decrease transmission, in the bottom panels, costs increase clearance rate. Other parameter values are $\beta = 3$, $\mu = 1$, $k_{MP} = k_{NM} = 0.5$, $k_{PN} = 1$ and all other $k = 0$

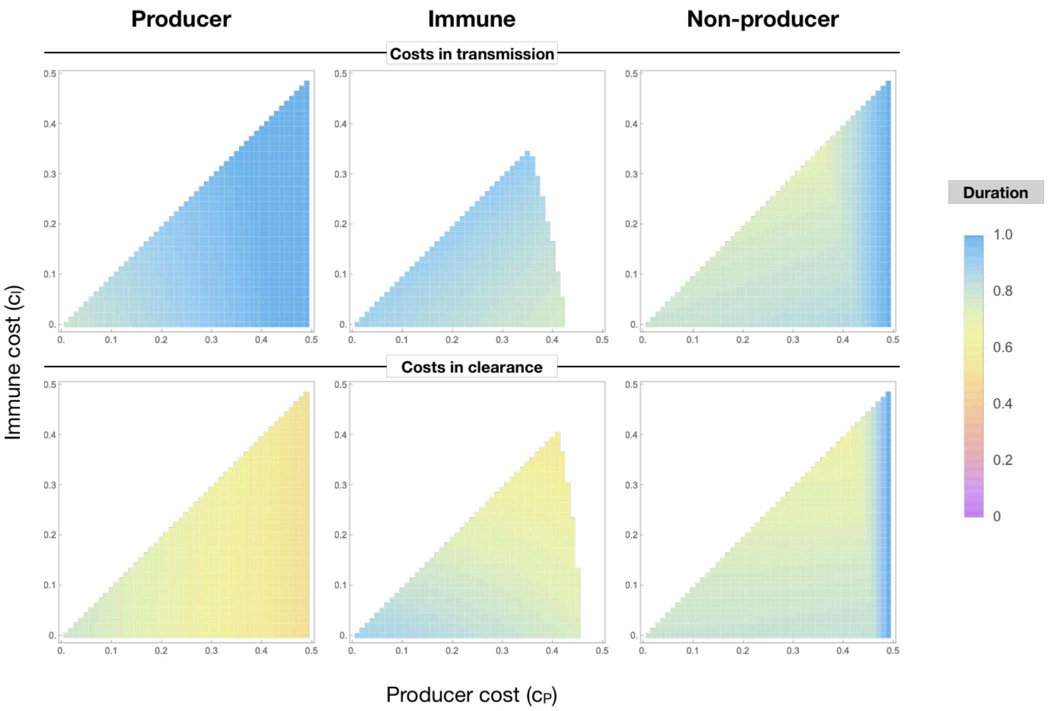


Figure S7: Duration of colonisation associated with each bacteriocin strain, for the same parameters as in SI Figure S5. As with the main text figure, for each panel, the x-axis represents the fitness cost incurred by the producer strain; the y-axis the fitness cost incurred by the immune strain. White space occurs when no duration is recorded. This happens when $c_M > c_P$ as we assume the producer strain incurs a higher cost than the immune strain, or when a particular strain is absent at equilibrium for particular values of fitness cost. In the top panels, fitness costs decrease transmission, in the bottom panels, costs increase clearance rate. Other parameter values are $\beta = 3$, $\mu = 1$, $k_{MP} = k_{NM} = 0.25$, $k_{PN} = 0.5$ and all other $k = 0$

#### 2.6 Resistance-associated cost in clearance

SI Figure S8 shows the effect of modelling the cost of resistance as affecting clearance rather than transmission. The effect in the main text (i.e. variation in duration of colonisation resulting in coexistence of antibiotic sensitive and resistant strains) is robust when both resistance- and bacteriocin-associated costs are modelled as affecting clearance rate. However, when bacteriocin-associated costs are modelled as affecting transmission rate and the cost of resistance is modelled as affecting only clearance rate ($\mu_R = \mu_S/(1-c_R)$), but not displacement ($k_{AB^R} = k_{AB^S}$, where $A$ and $B$ indicate strains), coexistence of sensitive and resistant strains is not observed. Coexistence is restored if the cost of resistance is also assumed to affect clearance through displacement: i.e. to increase the susceptibility of resistance strains to invasion ($k_{AB^R} = k_{AB^S}/(1-c_R)$).

To understand why, we consider a strain that can be cleared through two processes which occur at rates $\alpha$ and $\gamma$ and follow the reasoning in Lehtinen et al. [4]. If the cost of resistance affects transmission rate, the $R_0$ of the resistant strain is $\frac{(1-c_R)\beta}{\alpha+\gamma}$. The $R_0$ of the antibiotic sensitive strain is $\frac{\beta}{\alpha+\gamma+\tau}$, where $\tau$ is the population antibiotic consumption. The strain with the higher $R_0$ will out-compete the other—resistance is therefore selected for when $\tau > \frac{(\mu+\gamma)c_R}{1-c_R}$. If the cost is modelled in clearance and affects both components of clearance rate ($\alpha$ and $\gamma$), the expression for the $R_0$ of the resistant strain is identical and the result is the same. However, if only one of the processes (say the one occurring at rate $\alpha$) is affected by the cost of clearance, the $R_0$ of the resistant strain is $\frac{\beta}{\frac{\alpha}{(1-c_R)}+\gamma}$. Resistance is therefore selected for when $\tau > \frac{\mu c_R}{1-c_R}$.

Thus, if the cost of resistance affects clearance rate only, it is only the components of clearance rate affected by the cost of resistance that play a role in resistance dynamics. In the bacteriocin model, when bacteriocin-associated costs affect transmission, variation in duration of colonisation only arises from differences in susceptibility to displacement. Thus, variation in duration of colonisation only modulates the fitness of resistance when the cost of resistance affects susceptibility to displacement.

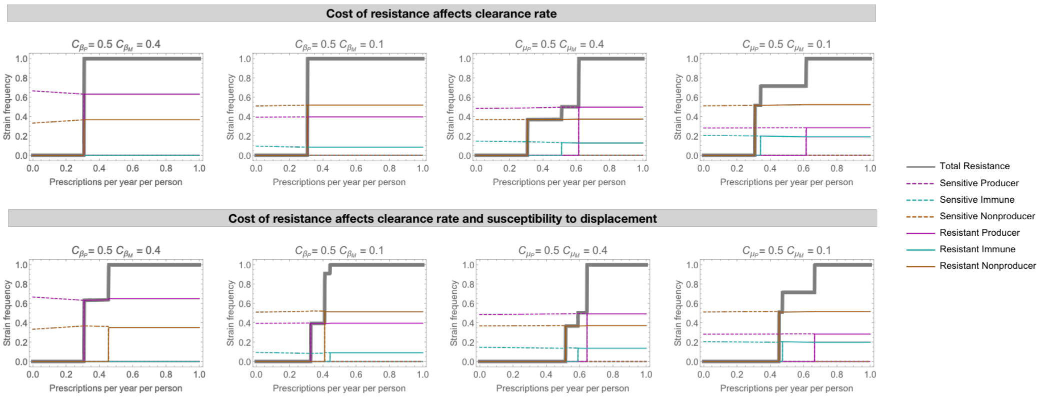


Figure S8: The effect of modelling the cost of antibiotic resistance as affecting clearance rather than transmission (as in the main text). In the top panels, the cost of resistance affects only the natural clearance rate $\mu$. In the bottom panels, the cost of resistance affects both natural clearance rate and susceptibility to replacement through invasion. As in the main text, strain frequencies (colours) and overall resistance frequency (gray) as a function of the population antibiotic consumption rate ($\tau$) for different parametrisations of the cost of the producer and immune strains. $c_\beta$ indicates cost in transmission and $c_\mu$ indicates cost in clearance, with $P$ and $M$ indicating the cost to the producer and immune strain respectively: in the top row, bacteriocin-associated costs affect transmission, in the bottom row these affect clearance. Other parameters are $\beta = 3$, $\mu = 1$, $k_{PN} = 1$, $k_{MP} = k_{NM} = 0.5$. Due to the difficulty of computing equilibrium solutions and eigenvalues for this model, these results are derived through simulating rather than stability analysis (simulation until $t = 10^5$ months).

### References


[1] Hjálmarsdóttir MÁ, Gumundsdóttir PF, Erlendsdóttir H, Kristinsson KG, Haraldsson G. Cocolonization of pneumococcal serotypes in healthy children attending day care centers: molecular versus conventional methods. The Pediatric infectious disease journal. 2016;35(5):477–480.

[2] Kamng'ona AW, Hinds J, Bar-Zeev N, Gould KA, Chaguza C, Msefula C, et al. High multiple carriage and emergence of Streptococcus pneumoniae vaccine serotype variants in Malawian children. BMC Infectious Diseases. 2015;15(1):234.

[3] Turner P, Hinds J, Turner C, Jankhot A, Gould K, Bentley SD, et al. Improved detection of nasopharyngeal cocolonization by multiple pneumococcal serotypes by use of latex agglutination or molecular serotyping by microarray. Journal of Clinical Microbiology. 2011;49(5):1784–1789.

[4] Lehtinen S, Blanquart F, Croucher NJ, Turner P, Lipsitch M, Fraser C. Evolution of antibiotic resistance is linked to any genetic mechanism affecting bacterial duration of carriage. Proceedings of the National Academy of Sciences of the United States of America. 2017 jan;114(5):1075–1080.
